# Ophiostomatalean fungi associated with bark beetles infesting *Pinus koraiensis*, including the description of *Ceratocystiopsis fushunensis* sp. nov. in China

**DOI:** 10.1101/2024.11.06.622286

**Authors:** Yutong Ran, Minjie Chen, Congwang Liu, Kun Liu, Tong Lin, Mingliang Yin

## Abstract

Fungi of Ophiostomatales (Sordariomycetes, Ascomycota) always have a symbiotic relationship with bark beetles (Coleoptera: Scolytinae). In this study, we investigated the fungi associated with *Cryphalus piceae* and *Hylastes* sp in Fushun. Based on the four gene fragments of TUB, ITS, LSU, and TEF and morphological characteristics, 206 strains of Ophiostomatales were identified to be distributed in 21 taxa; these included an undescribed taxon. Here, we described it as *Ceratocystiopsis fushunensis* sp. nov. The results of this study provided valuable information on the relationship between fungi and bark beetles.

## 1. Introduction

Ophiostomatales was first proposed by Benny & Kimbrough (1981), including one family, Ophiostomataceae; four genera, *Ophiostoma, Ceratocystiopsis, Sphaeronaemella* and *Ceratocystis*. Hausner et al. (1993) proposed the transfer of *Ceratocystiopsis* into *Ophiostoma* based on the characterization of ascospores, supported by Viljoen et al. (2000) with a detailed description of morphological characterization. *Grosmannia* and *Ceratocystiopsis* were restored by Zipfel et al. (2006) using DNA sequence analysis of partial LSU and TUB genes. De Beer et al. (2013a) redefined *Ceratocystiopsis, Fragosphaeria, Graphilbum*, etc., and identified 18 complexes of different genera. After that, scholars focused on the reports of new species and the revision of the complex (Feau et al., 2024; Linnakoski et al., 2016; Musvuugwa et al., 2015; Simmons et al., 2016; Wang et al., 2022; Yin et al., 2020). De Beer et al. (2022) amplified and sequenced the DNA of more than 200 species, redefined Ophiostomatales, described all genera in detail, and provided the most comprehensive phylogenetic data of Ophiostomatales. Ophiostomatales currently includes one family, 16 genera, and about 800 species.

Ophiostomatales fungi are a kind of fungi associated with bark beetles, which can adapt well to the spread of arthropods (Malloch & Blackwell, 1992). These fungi are adapted for transmission by bark beetles or other insects by producing large numbers of sticky ascospores to ensure they are not easily removed during transportation (Upadhyay, 1981). Ophiostomatales fungi act as a beneficial resource as food for beetles (Lewis & Alexander, 1986) or play a role in the development of larvae (Brand et al., 1976). In addition, studies have also shown that *Hylastes sclerosis* and *H. tenuis* reproduction rates are increased by Ophiostomatales fungi (Eckhardt et al., 2004), which in turn help the beetle overcome tree defenses by detoxifying plant antitoxins (Raffa et al., 2015; Wadke et al., 2016).

*Hylastes* Erichson (Coleoptera: Scolytinae) mainly feeds on the phloem and cambium of dead or dying pine trees. It exists in the stumps and roots of pine trees (Schedl, 1963; Schwenke, 1982), infesting mainly *Pinus elliottii, P. laricio, P. sylvestris, Larix decidua* and other tree species (Bright, 2014; Paine & Lieutier, 2016).

Some *Hylastes* spp. Invade vulnerable pines and spread *Grosmannia alacris, Leptographium terebrantis*, and their interactions can further lead to pine decline (Eckhardt, 2002; Zeng et al., 2014), and again *Hylastes porculus* Erichson is the primary vector of *Pine koraiensis* root disease caused by the blue-stain fungus *Leptographium procerum* (Klepzig et al., 1991; Schowalter, 2018). Zhou et al. (2001) isolated seven species of Ophiostomatales fungi, including *Ophiostoma ips, Ceratocystiopsis minuta, Sporothrix* sp., and *Leptogmphium lundbergii*, from *H. angustatus* and its galleries. *Cryphalus piceae* (Coleoptera: Scolytinae) is the common species of *Cryphalus* Erichson, which mainly infects *Pinus, Picea*, and *Larix* in Europe and Asia (Cerchiarini et al., 1997). It is a secondary bark beetle that primarily attacks dying trunks of trees no less than 8-10 years old, as well as fallen branches (Justesen et al., 2020). Scholars have isolated fungi of several genera including *Sporothrix, Graphilbum, Lepptographium, Ophiostoma, Grosmannia*, and others from *C. piceae* infested pine trees in Poland and Japan. At present, more than 60 species of fungi related to this bark beetle have been found (Jankowiak & Bilański, 2018; Jankowiak & Kolařík, 2010; Jankowiak et al. 2017; Ohtaka et al. 2002a; Ohtaka et al. 2002b).

Liaoning Province is located at the intersection of Changbai Mountain, North China, and Inner Mongolia flora, with rich forest resources that provide a good breeding ground for insects and fungal organisms. In August 2022, we sampled the blue-stain wood, bark beetles, and their galleries in Dalian and Fushun and isolated 206 strains of Ophiostomatales. Through morphological observation and multi-locus DNA sequence data analysis, 21 species of Ophiostomatales fungi in 6 genera were obtained. Combined with phylogenetic analysis, a new species was identified. Here, we describe it as *Ceratocystiopsis fushunensis* sp. nov. The results of this study enriched the diversity of Ophiostomatales fungi resources and provided a basis for further understanding the relationship between Ophiostomatales fungi and bark beetles in the Liaoning region.

## 2. Methods and materials

### 2.2 Sample collection and isolation

In August 2022, beetles and their galleries were collected from Dalian (38° 92′ N, 121° 62′ E) and Fushun (41° 52′ N, 123° 57′ E), Liaoning Province. The galleries collected in this study were all taken from dead trees that showed signs of wilting or dying due to the damage of bark beetles. The beetles were placed separately in 1.5 mL Eppendorf tubes, and the galleries and blue-stain wood were sealed with envelope bags. Before fungal isolation, they were stored at 4°C. Fungi isolated from bark beetles or galleries were inoculated onto 90 mm of 2% malt extract agar medium (20g Biolab agar and 1,000 mL deionized water) and placed in an artificial climatic chamber at 25°C for 7 days. The mycelial apex purification method was then used to purify the fungi cultured from the tissue isolates, which were inoculated on a 2% MEA medium and placed in an artificial climate chamber for dark culture under the same conditions. All fungal isolates in this study were preserved in South China Agricultural University, Guangzhou, China. Ex-holotype cultures of Ophiostomatales fungi described in this study were deposited in the China General Microbiological Culture Collection Center (CGMCC; http://www.cgmcc.net/english/catalogue.html), Beijing, China. Holotype specimens (dry cultures) were deposited in the Herbarium Mycologicum, Academiae Sinicae (HMAS), Beijing, China.

### 2.2 DNA extraction, PCR amplification and sequencing

DNA extraction was performed on the purified strains cultured in the dark for 7 days. The active aerial hyphae on the surface of the MEA plate were picked into 1.5 mL sterile Eppendorf tubes. DNA was extracted using PrepMan™ Ultra DNA Extraction Reagent (by Thermo Fisher Scientific) according to the instruction manual, and the extracted DNA stock solution was placed at four °C for storage.

Four gene fragments were amplified and sequenced using four pairs of primers. Bt2a/Bt2b primers (Glass & Donaldson, 1995) were used to amplify partial genes β-tubulin (TUB). ITS5/ITS4 primers (White et al., 1990) were used to amplify the internal transcribed spacer (ITS). LROR/LR5 primers (Vilgalys & Hester, 1990) were used to amplify the ribosomal large subunit region (LSU). And EF1F/EF2R (Jacobs et al., 2004) or EF2F (Marincowitz et al., 2015)/EF2R were used to amplify the translation elongation factor 1-α (EF1-α) gene region (TEF).

A 25uL PCR reaction system was used. The PCR reaction system included 12.5uL 2 × Taq Master Mix (by Thermo Fisher Scientific), 0.5uL forward system primers, 0.5uL reverse system primers, 10.5uL PCR grade deionized water, and 1uL DNA stock solution. The amplification conditions were set as pre-denaturation at 95°C for 3 minutes, denaturation at 95°C for 30 seconds, annealing at 55°C for 30 seconds, extension at 72°C for 1 minute, 35 cycles, and then final extension at 72°C for 10 minutes, stored at 12°C. The annealing temperature varied between 52°C-58°C depending on the primers used. The successfully amplified PCR samples were sent to Sangon Biotech (Shanghai) Co., Ltd. for sequencing.

### 2.3 Phylogenetic analysis

After successful sequencing, the obtained TUB gene sequences were searched in the NCBI Genbank database, and the target sequence was subjected to gene sequence alignment and homologous similarity search using BLAST to determine the genus and species of the sequenced strain initially. The sequences of the four gene regions of TUB, ITS, LSU, and TEF of the experimental strains in this study were organized, and representative sequences, model strain sequences, and related species sequences with high matching degrees were downloaded. The sequences of the four gene regions were organized in MAFFT v. 7 (https://mafft.cbrc.jp/alignment/server/) (Katoh & Standley, 2013) using the E-INS-i strategy for sequence alignment, followed by manual editing of the sequences on the software MEGA 7.0 (Kumar et al., 2016). For maximum likelihood (ML) and Bayesian inference (BI) analyses, the best alternative model was selected for each data set at jModelTest 2.1.10 (Darriba et al., 2012). Processed data were used to construct phylogenetic trees (1000 bootstrap pseudoreplicates) with ML on MEGA X; Maximum parsimony (MP) analysis was performed using PAUP version 4.0b10 (Swofford et al., 2002), with gaps as the fifth base; and the data were analyzed using MrBayes v3.2.7 (Ronquist et al., 2012) for BI analysis, with five million generations run simultaneously on the four MCMC chains by the Markov Chain Monte Carlo (MCMC) method. Phylogenetic trees were edited visually with Figtree v.1.4.3.

### 2.4 Morphological and Physiological Characteristics

Morphological features were observed and recorded using a stereoscope (Leica M165C), optical microscope (OLYMPUS-BX43), and camera (Nikon). To determine the growth rate of the new species of fungi, the active fungal colonies on the isolated bacteria were selected, and the agar blocks with a diameter of 5mm were inoculated into a 2% MEA culture dish with a diameter of 90mm, and cultured in darkness at 25°C. The diameter of the two vertical axes was measured on days 7 and 14 until the hyphae covered the edge of the MEA culture dish. At the same time, the color of the surface and back of the colony was judged according to Rayner’s chromaticity diagram (Rayner, 1970).

The asexual or sexual morphology of fungi was induced by pine strip plate to observe the microstructure of fungi. The seedlings were inoculated on a horizontal plate of pine strips and cultured in a dark environment at 25°C. The growth in the plate was observed regularly by a stereoscope. After 14 days, the mycelium was picked up to make slides, and the microstructure photos were observed and collected under the optical microscope. At the same time, 30 measurements were repeated for each reproductive structure of the strain, expressed as (min-) (mean-SD) - (mean + SD) (-max). The data of all strains were stored in MycoBank (http://www.MycoBank.org/).

## 3. Results

### 3.1 Sample collection and isolation

In this study, 448 bark beetles were collected from *P. koraiensis*, 402 from *Hylastes* sp., and 46 from *C. piceae*. Two hundred six strains of Ophiostomatales fungi were isolated from the two species of bark beetles. Of these, 183 strains of fungi were collected from *Hylastes* sp. and its galleries, and 23 strains of fungi were collected from *C. piceae* and galleries.

### 3.2 Phylogenetic analysis

The BLAST alignment of the TUB sequences of the strains isolated in this study showed that 226 strains of Ophiostomatales fungi belonged to 6 genera, namely *Sporothrix* (17 strains), *Ophiostoma* (29 strains), *Leptographium* (65 strains), *Graphilbum* (70 strains), *Ceratocystiopsis* (17 strains), *Masuyamyces* (8 strains). Analysis of the ML tree constructed based on TUB sequences (Fig. 1) showed that these strains were distributed in 21 taxa, of which 20 were known taxon and one was unknown.

**Fig. 1.**
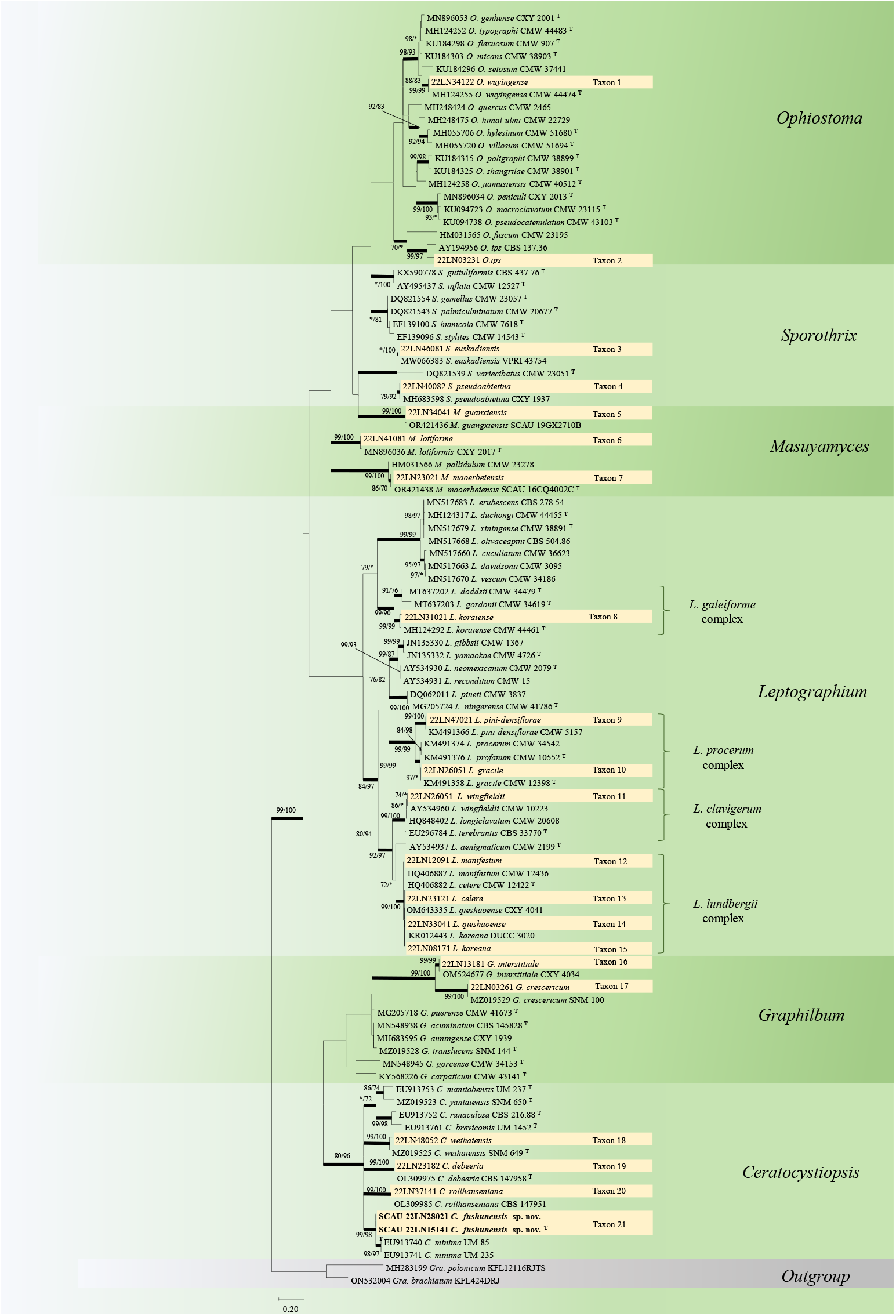
Phylogram obtained from ML analyses of the TUB data for six genera of Ophiostomatales. Bootstrap values ≥ 70% for ML and MP analyses are presented at nodes as follows: ML/MP. Bold branches indicate posterior probabilities values ≥ 0.95 obtained from BI analyses. *Bootstrap values < 70%. T = ex-type cultures.

The sequences related to *Ceratocystiopsis* were collected (Table 1), and phylogenetic analysis of the *Ceratocystiopsis* was performed based on the TUB, ITS, TEF, and TUB-ITS-LSU combined fragments. ML, MP, and BI processed four fragments, the best model for TUB, ITS, TUB-ITS-LSU fragment were GTR+I+G, and the best model for TEF was TrN+I+G (Table 2). The results showed that in the TUB, ITS trees (Fig. 2), the four strains of Taxon 21 were clustered into a single evolutionary branch with high node support, closely linked to strain *C. minima* (UM 85, UM 235). In the TEF, TUB-ITS-LSU trees (Fig. 3), Taxon 21 was separated from the other strains with full support, closely connected to strain *C. yantaiensis* (SNM 650, SNM 582) of the TEF tree. The phylogenetic trees showed that the four strains were grouped into a separate lineage, and all four phylogenetic trees demonstrated that Taxon1 is a new species that has never been described.

**Table 1.**
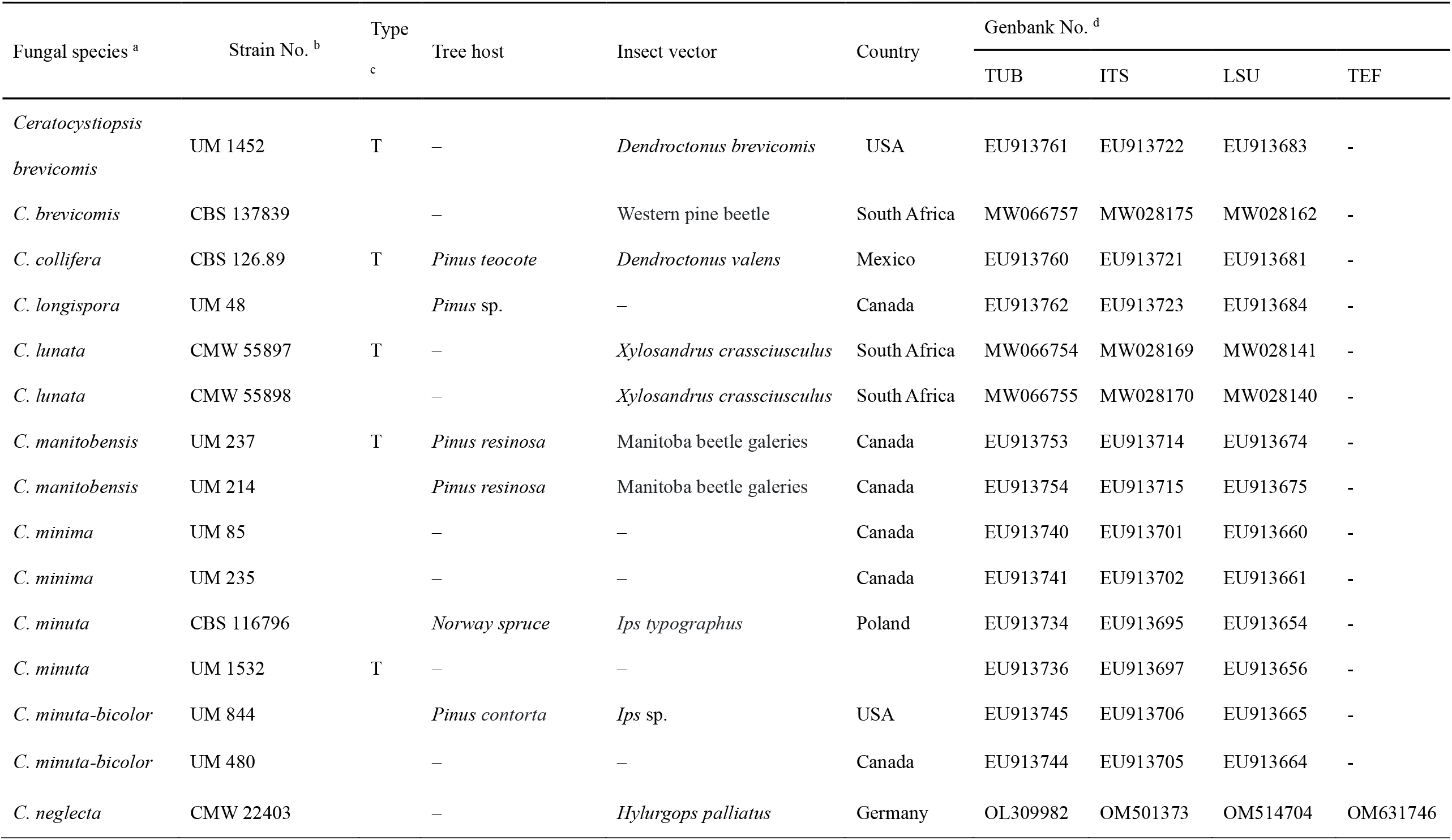
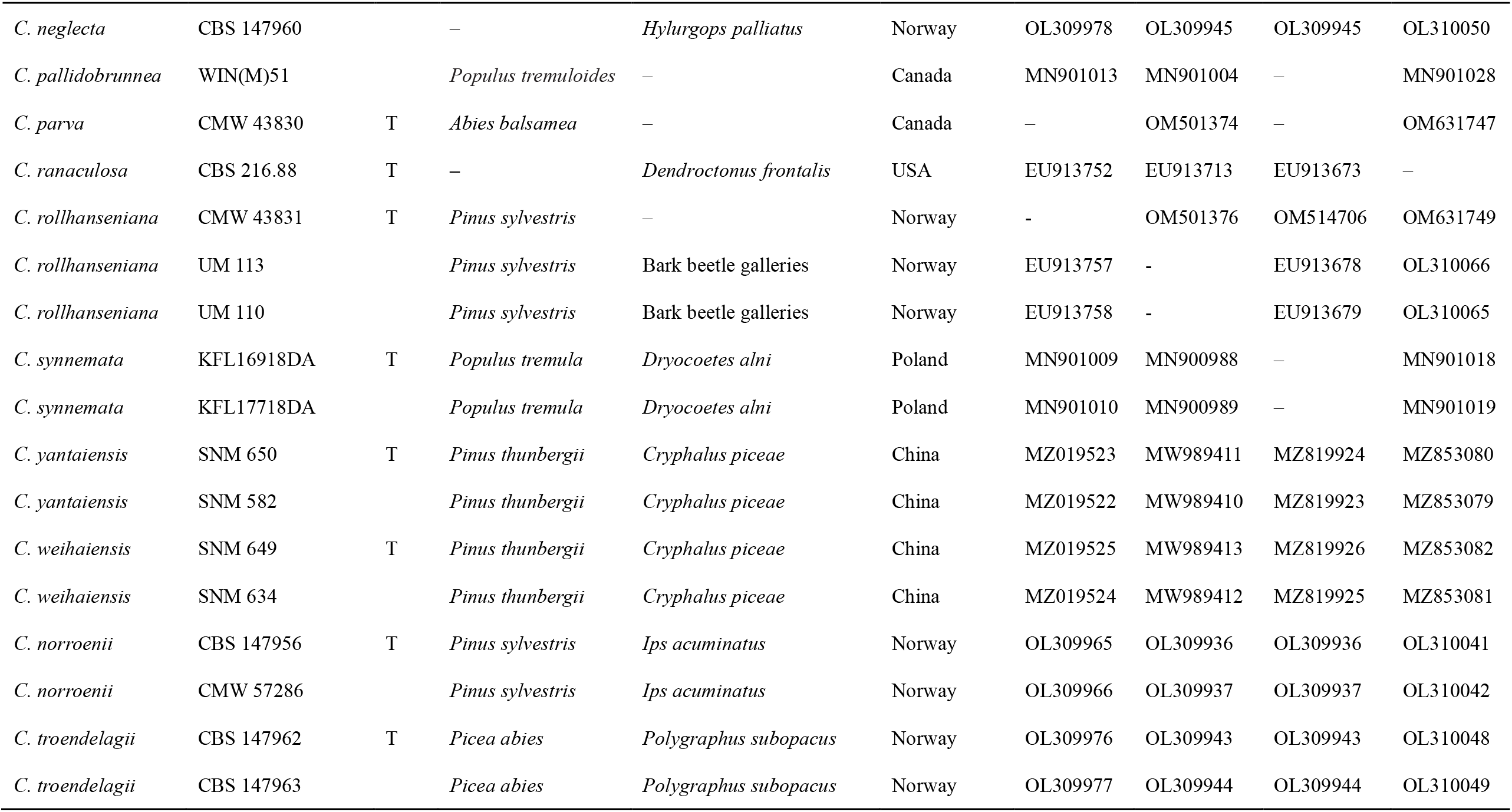
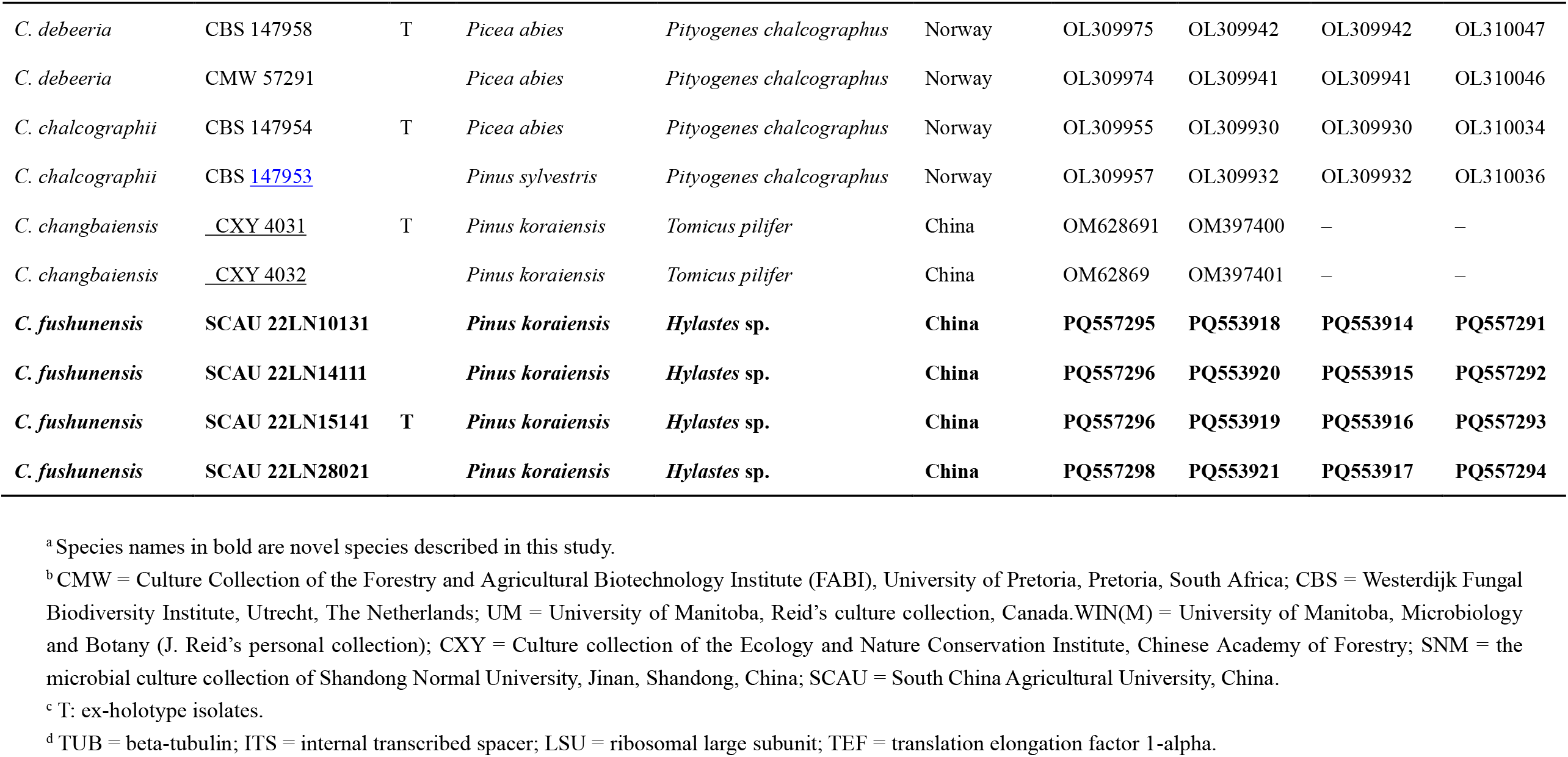
Representative strains used for phylogenetic analysis in this study.

| Fungal species <sup>a</sup> | Strain No. <sup>b</sup> | Type <sup>c</sup> | Tree host | Insect vector | Country | Genbank No. <sup>d</sup> |  |  |  |
| --- | --- | --- | --- | --- | --- | --- | --- | --- | --- |
|  |  |  |  |  |  | TUB | ITS | LSU | TEF |
| *Ceratocystiopsis brevicomis* | UM 1452 | T | – | *Dendroctonus brevicomis* | USA | EU913761 | EU913722 | EU913683 | - |
| *C. brevicomis* | CBS 137839 |  | – | Western pine beetle | South Africa | MW066757 | MW028175 | MW028162 | - |
| *C. collifera* | CBS 126.89 | T | *Pinus teocote* | *Dendroctonus valens* | Mexico | EU913760 | EU913721 | EU913681 | - |
| *C. longispora* | UM 48 |  | *Pinus* sp. | – | Canada | EU913762 | EU913723 | EU913684 | - |
| *C. lunata* | CMW 55897 | T | – | *Xylosandrus crassiusculus* | South Africa | MW066754 | MW028169 | MW028141 | - |
| *C. lunata* | CMW 55898 |  | – | *Xylosandrus crassiusculus* | South Africa | MW066755 | MW028170 | MW028140 | - |
| *C. manitobensis* | UM 237 | T | *Pinus resinosa* | Manitoba beetle galleries | Canada | EU913753 | EU913714 | EU913674 | - |
| *C. manitobensis* | UM 214 |  | *Pinus resinosa* | Manitoba beetle galleries | Canada | EU913754 | EU913715 | EU913675 | - |
| *C. minima* | UM 85 |  | – | – | Canada | EU913740 | EU913701 | EU913660 | - |
| *C. minima* | UM 235 |  | – | – | Canada | EU913741 | EU913702 | EU913661 | - |
| *C. minuta* | CBS 116796 |  | *Norway spruce* | *Ips typographus* | Poland | EU913734 | EU913695 | EU913654 | - |
| *C. minuta* | UM 1532 | T | – | – |  | EU913736 | EU913697 | EU913656 | - |
| *C. minuta-bicolor* | UM 844 |  | *Pinus contorta* | *Ips* sp. | USA | EU913745 | EU913706 | EU913665 | - |
| *C. minuta-bicolor* | UM 480 |  | – | – | Canada | EU913744 | EU913705 | EU913664 | - |
| *C. neglecta* | CMW 22403 |  | – | *Hylurgops palliatus* | Germany | OL309982 | OM501373 | OM514704 | OM631746 |

| Fungal species <sup>a</sup> | Strain No. <sup>b</sup> | Type<br><sup>c</sup> | Tree host | Insect vector | Country | Genbank No. <sup>d</sup> |  |  |  |
| --- | --- | --- | --- | --- | --- | --- | --- | --- | --- |
|  |  |  |  |  |  | TUB | ITS | LSU | TEF |
| *C. neglecta* | CBS 147960 |  | – | *Hylurgops palliatus* | Norway | OL309978 | OL309945 | OL309945 | OL310050 |
| *C. pallidobrunnea* | WIN(M)51 |  | *Populus tremuloides* | – | Canada | MN901013 | MN901004 | – | MN901028 |
| *C. parva* | CMW 43830 | T | *Abies balsamea* | – | Canada | – | OM501374 | – | OM631747 |
| *C. ranaculosa* | CBS 216.88 | T | – | *Dendroctonus frontalis* | USA | EU913752 | EU913713 | EU913673 | – |
| *C. rollhanseniana* | CMW 43831 | T | *Pinus sylvestris* | – | Norway | - | OM501376 | OM514706 | OM631749 |
| *C. rollhanseniana* | UM 113 |  | *Pinus sylvestris* | Bark beetle galleries | Norway | EU913757 | - | EU913678 | OL310066 |
| *C. rollhanseniana* | UM 110 |  | *Pinus sylvestris* | Bark beetle galleries | Norway | EU913758 | - | EU913679 | OL310065 |
| *C. synnemata* | KFL16918DA | T | *Populus tremula* | *Dryocoetes alni* | Poland | MN901009 | MN900988 | – | MN901018 |
| *C. synnemata* | KFL17718DA |  | *Populus tremula* | *Dryocoetes alni* | Poland | MN901010 | MN900989 | – | MN901019 |
| *C. yantaiensis* | SNM 650 | T | *Pinus thunbergii* | *Cryphalus piceae* | China | MZ019523 | MW989411 | MZ819924 | MZ853080 |
| *C. yantaiensis* | SNM 582 |  | *Pinus thunbergii* | *Cryphalus piceae* | China | MZ019522 | MW989410 | MZ819923 | MZ853079 |
| *C. weihaiensis* | SNM 649 | T | *Pinus thunbergii* | *Cryphalus piceae* | China | MZ019525 | MW989413 | MZ819926 | MZ853082 |
| *C. weihaiensis* | SNM 634 |  | *Pinus thunbergii* | *Cryphalus piceae* | China | MZ019524 | MW989412 | MZ819925 | MZ853081 |
| *C. norroenii* | CBS 147956 | T | *Pinus sylvestris* | *Ips acuminatus* | Norway | OL309965 | OL309936 | OL309936 | OL310041 |
| *C. norroenii* | CMW 57286 |  | *Pinus sylvestris* | *Ips acuminatus* | Norway | OL309966 | OL309937 | OL309937 | OL310042 |
| *C. troendelagii* | CBS 147962 | T | *Picea abies* | *Polygraphus subopacus* | Norway | OL309976 | OL309943 | OL309943 | OL310048 |
| *C. troendelagii* | CBS 147963 |  | *Picea abies* | *Polygraphus subopacus* | Norway | OL309977 | OL309944 | OL309944 | OL310049 |
|  |  |  |  |  |  | TUB | ITS | LSU | TEF |
| *C. debeeria* | CBS 147958 | T | *Picea abies* | *Pityogenes chalcographus* | Norway | OL309975 | OL309942 | OL309942 | OL310047 |
| *C. debeeria* | CMW 57291 |  | *Picea abies* | *Pityogenes chalcographus* | Norway | OL309974 | OL309941 | OL309941 | OL310046 |
| *C. chalcographii* | CBS 147954 | T | *Picea abies* | *Pityogenes chalcographus* | Norway | OL309955 | OL309930 | OL309930 | OL310034 |
| *C. chalcographii* | CBS 147953 |  | *Pinus sylvestris* | *Pityogenes chalcographus* | Norway | OL309957 | OL309932 | OL309932 | OL310036 |
| *C. changbaiensis* | <u>CXY 4031</u> | T | *Pinus koraiensis* | *Tomicus pilifer* | China | OM628691 | OM397400 | – | – |
| *C. changbaiensis* | <u>CXY 4032</u> |  | *Pinus koraiensis* | *Tomicus pilifer* | China | OM62869 | OM397401 | – | – |
| ***C. fushunensis*** | **SCAU 22LN10131** |  | ***Pinus koraiensis*** | ***Hylastes* sp.** | **China** | **PQ557295** | **PQ553918** | **PQ553914** | **PQ557291** |
| ***C. fushunensis*** | **SCAU 22LN14111** |  | ***Pinus koraiensis*** | ***Hylastes* sp.** | **China** | **PQ557296** | **PQ553920** | **PQ553915** | **PQ557292** |
| ***C. fushunensis*** | **SCAU 22LN15141** | T | ***Pinus koraiensis*** | ***Hylastes* sp.** | **China** | **PQ557296** | **PQ553919** | **PQ553916** | **PQ557293** |
| ***C. fushunensis*** | **SCAU 22LN28021** |  | ***Pinus koraiensis*** | ***Hylastes* sp.** | **China** | **PQ557298** | **PQ553921** | **PQ553917** | **PQ557294** |

<sup>a</sup> Species names in bold are novel species described in this study.
<sup>b</sup> CMW = Culture Collection of the Forestry and Agricultural Biotechnology Institute (FABI), University of Pretoria, Pretoria, South Africa; CBS = Westerdijk Fungal Biodiversity Institute, Utrecht, The Netherlands; UM = University of Manitoba, Reid's culture collection, Canada. WIN(M) = University of Manitoba, Microbiology and Botany (J. Reid's personal collection); CXY = Culture collection of the Ecology and Nature Conservation Institute, Chinese Academy of Forestry; SNM = the microbial culture collection of Shandong Normal University, Jinan, Shandong, China; SCAU = South China Agricultural University, China.
<sup>c</sup> T: ex-holotype isolates.
<sup>d</sup> TUB = beta-tubulin; ITS = internal transcribed spacer; LSU = ribosomal large subunit; TEF = translation elongation factor 1-alpha.

**Table 2.** Substitutional models data used for phylogenetic analysis in this study.

| Dataset | Maximum Likelihood |  |  | Maximum Parsimony |  |  |  |  |  | Bayesian inference |  |
| --- | --- | --- | --- | --- | --- | --- | --- | --- | --- | --- | --- |
|  | Subst.model | P-inv | Gamma | No.of trees | Tree length | CI | RI | RC | HI | Subst.model | Burn-in |
| TUB(Fig. 1) | GTR+I+G | 0.314 | 1.200 | 335 | 1936 | 0.427 | 0.845 | 0.361 | 0.573 | GTR+I+G | 1.25×10<sup>6</sup> |
| TUB(Fig. 2) | GTR+I+G | 0.402 | 1.219 | 1 | 996 | 0.532 | 0.814 | 0.433 | 0.467 | GTR+I+G | 1.25×10<sup>6</sup> |
| ITS | GTR+I+G | 0.251 | 0.475 | 2 | 990 | 0.646 | 0.803 | 0.519 | 0.353 | GTR+I+G | 1.25×10<sup>6</sup> |
| TEF | TrN+I+G | 0.244 | 1.271 | 1 | 1492 | 0.651 | 0.784 | 0.510 | 0.349 | TrN+I+G | 1.25×10<sup>6</sup> |
| TUB-ITS-LSU | GTR+I+G | 0.413 | 0.468 | 3 | 1590 | 0.672 | 0.834 | 0.561 | 0.327 | GTR+I+G | 1.25×10<sup>6</sup> |

Note: Subst. Model = substitution models used in phylogenetic analyses; P-inv = proportion of invariable sites; Gamma = Gamma distribution shape parameter; CI = consistency index; RI = retention index; RC = rescaled consistency index; HI = homoplasy index.

**Fig. 2.**
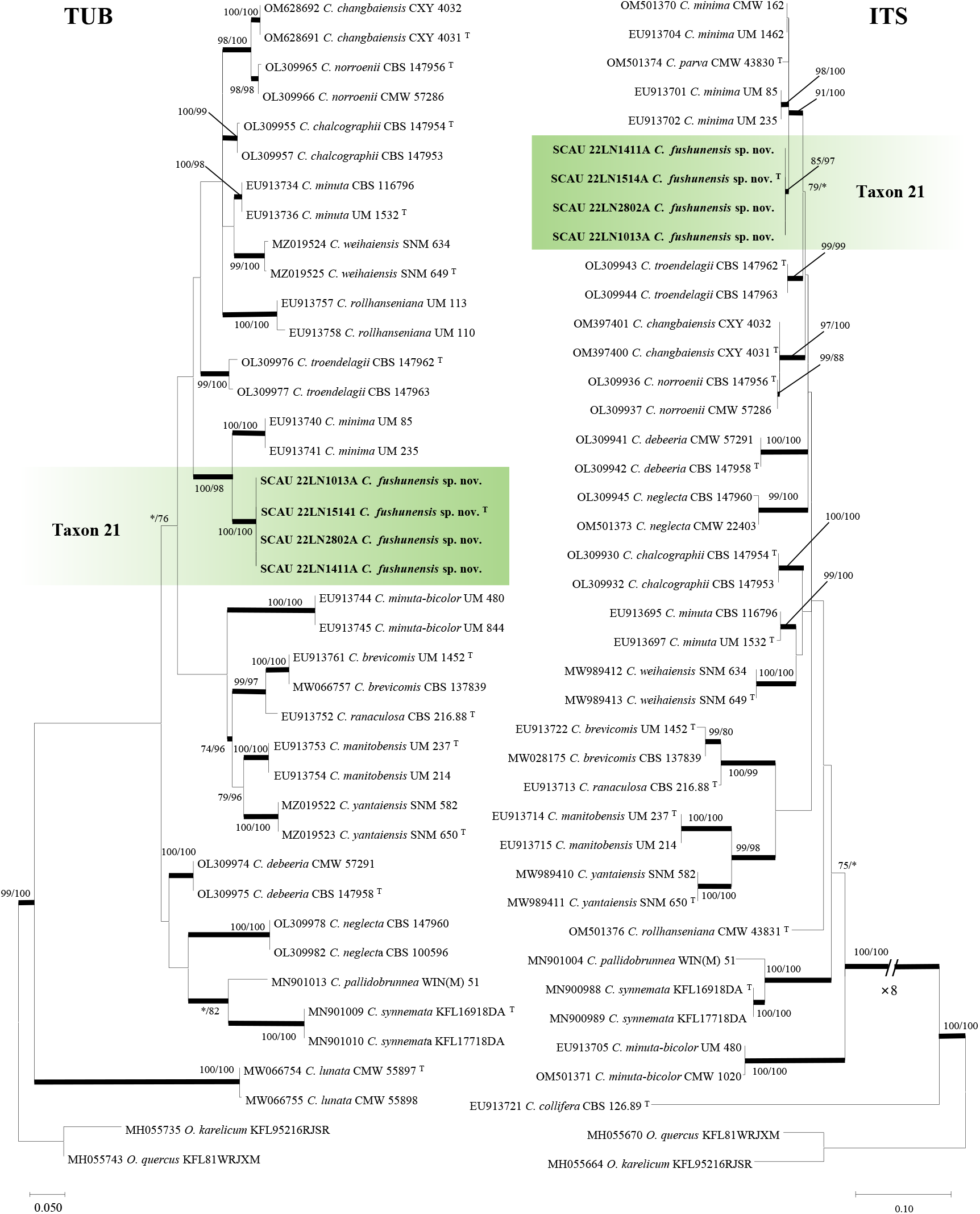
Phylogram obtained from ML analyses of the TUB and ITS data for *Ceratocystiopsis* spp. Bootstrap values ≥ 70% for ML and MP analyses are presented at nodes as follows: ML/MP. Bold branches indicate posterior probabilities values ≥ 0.95 obtained from BI analyses. *Bootstrap values < 70%. T = ex-type cultures.

**Fig. 3.**
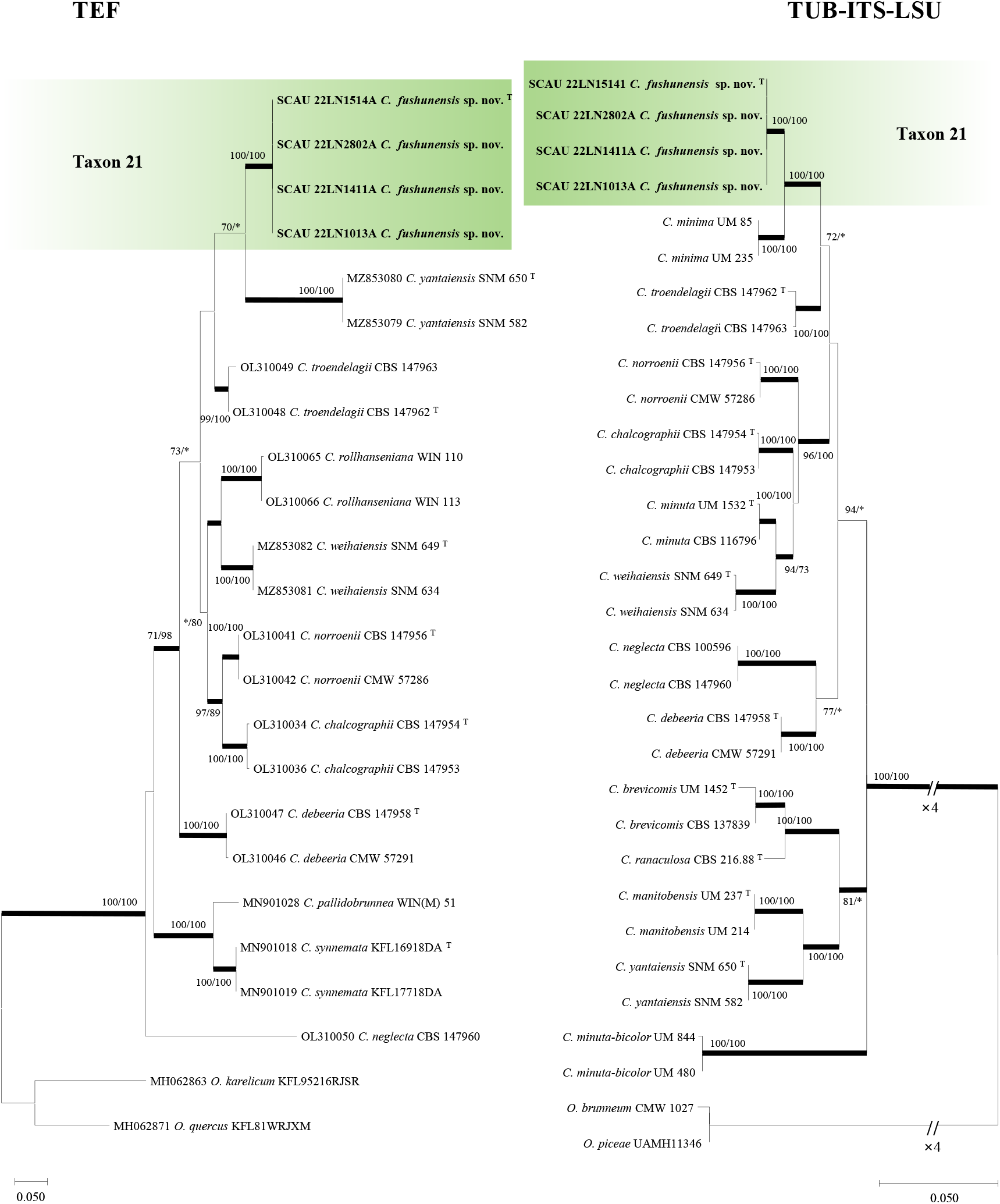
Phylogram obtained from ML analyses of the TEF and TUB-ITS-LSU data for *Ceratocystiopsis* spp. Bootstrap values ≥ 70% for ML and MP analyses are presented at nodes as follows: ML/MP. Bold branches indicate posterior probabilities values ≥ 0.95 obtained from BI analyses. *Bootstrap values < 70%. T = ex-type cultures.

### 3.3 Taxonomy

Comparisons based on morphological characters and multi-locus phylogenetic analyses indicate that Taxon 21 is distinct from other known taxon in Ophiostomatales. Therefore, it is described as a new species in this study:

Name: *Ceratocystiopsis fushunensis* Y.T. Ran & M.L. Yin, sp. nov. fig. 4.

**Fig. 4.**
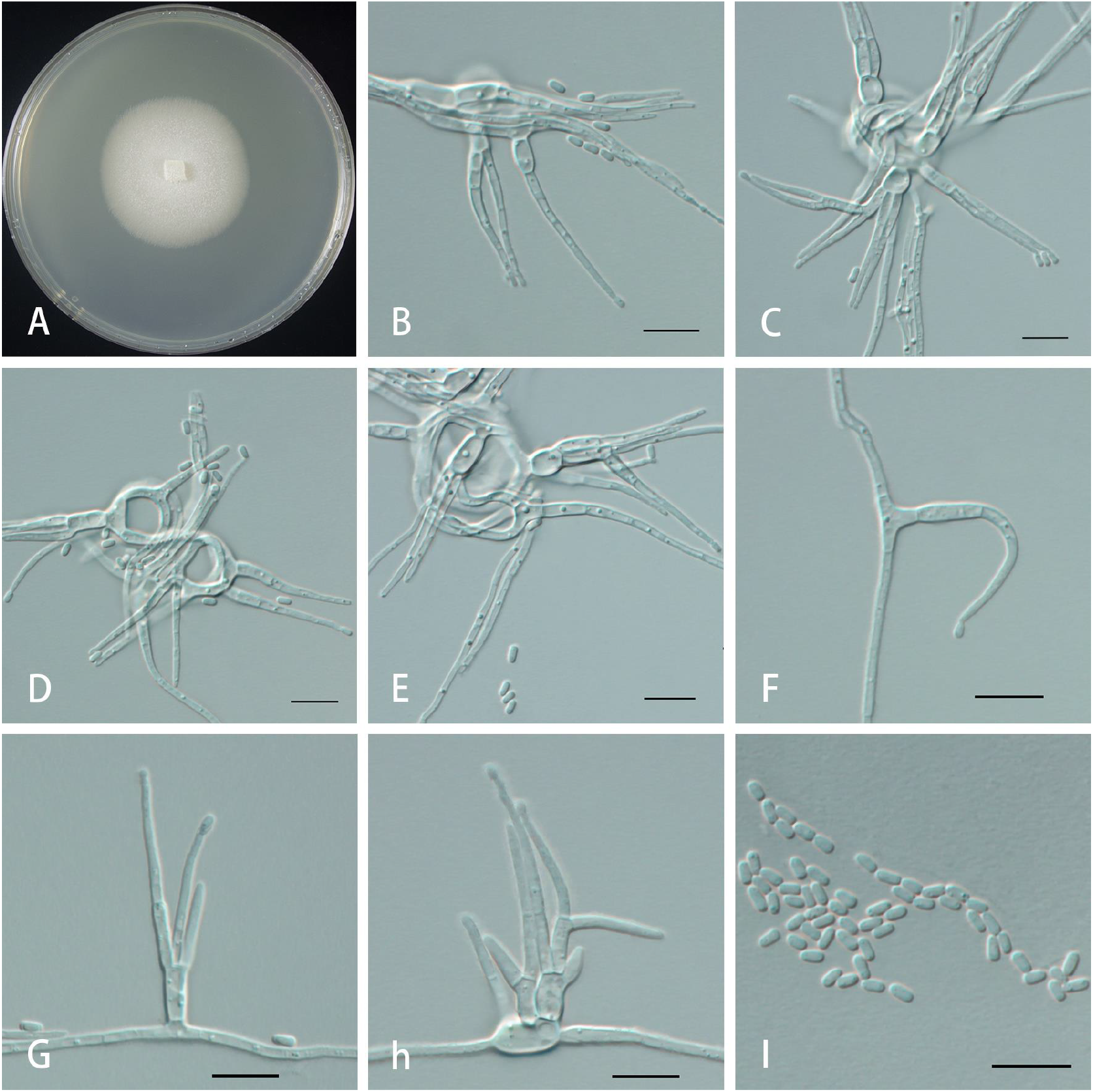
Morphological characteristics of asexual structures of *C. fushunensis sp. nov*.. **A**: Fourteen-day-old culture on MEA. **B-H**: Hyalorhinocladiella-like conidiophores. **I**: Conidia. Scale bar = 5 um.

MycoBank no.: MB 856473.

Type: China, Liaoning Province, Fushun City, taken from *P. koraiensis* infested with *Hylastes* sp., 4 Aug 2022. holotype HMAS 352763, ex-holotype CGMCC 3.27260 = SCAU 22LN15141.

Etymology: the etymology refers to the city of Fushun, where the fungus was isolated from a *P. koraiensis* sample.

Diagnosis: The sister species of this species is *C. minima*.

Description: No sexual form observed. The asexual form is hyalorhinocladiella-like. Linear conidiophores are produced by trophic mycelium. The conidiophores are simple, hyaline, branched, erect or curved. Conidiophores are (8.5-)11.1-20.9(−28.9) μm long and (0.9-)1-1.8(−2.7) μm wide at the base. Conidiophores are often 2-3-branched. Conidiophores are hyaline, smooth, unicellular, septate, ellipsoidal, or obovate, and slightly narrower at the base, (1.2-)1.3-1.7(−2.0) × (0.5-)0.6-(−0.9). 0.8(−0.9) μm.

Culture characteristics: On 2% MEA, the colony is white with a light orange color on the back. The aerial mycelium is fluffy and attached to the surface of the medium. The edge of the colony was radially thinning in the late stage of growth. The radial growth rate was 2.7 (±0.1) mm/d. The optimum growth temperature was 25°C, and no growth was below 5 °C or above 35°C.

Host trees: *P. koraiensis*

Insect vector: *Hylastes* sp.

Distribution: Lingning Province, China

Notes: According to phylogenetic analysis, Taxon1 is most closely related to *C*.*minima*, but there are significant differences in morphology. The latter was isolated from *P. banksiana* in Canada produces a sexual morphology with fusiform ascospores, with conidial peduncles up to 100 μm long, unbranched or hairy branched; there are also significant differences in conidial morphology, with *C. minima* conidia constricted in the middle, rod-shaped or oblong, 2.5-5.0 (8.0) × 0.7-1.5 (2.0) μm. The spore sizes of the two differed considerably.

## 4. Discussion

In this study, the barks of *P. koraiensis* infected by *C. piceae* and *Hylastes* sp. were collected, and 226 strains of fungi were isolated from the galleries. These were identified as 21 species of 6 genera, including a new species.

Our results identified a new species, bringing the total *Ceratocystiopsis* species to 35. The new species’ morphological characteristics are similar to those of other species of Ceratocystiopsis, with a hyalorhinocladiella-like asexual morphology (Beer et al., 2013), but there are differences in the size and morphology of conidia. Combined with phylogenetic analysis and morphological characteristics, it is easy to distinguish from other known species.

In our study, the dominant species is *Graphilbum interstitiale* (67 of 226 strains), accounting for 32.5 %. *G. interstitiale* was first isolated as a new species from *P. sylvestris* infected by *Hylurgops interstitialis* in the Russian Far East (Jankowiak et al., 2020) and was later found in *P. koraiensis* infected by *Tomicus pilifer* in Northeast China (Wang et al., 2022). *G. interstitiale* has not been reported to be associated with *Hylastes* sp., but this was found in our study. Three strains of *G. crescericum* in the genus were also isolated from the galleries of *C. piceae* in this study, accounting for 1.5% of the total, and its geographic distribution and hosts were the same as those previously found by Chang et al. (2021).

The data show a high diversity of fungi in the *Leptographium* (65 out of 226), containing eight species from four complexes in the genus, with 36 strains (17.4%) in the *L. lundbergii* complex, 27 strains (13.1%) in the *L. procerum* complex, *L. clavigerum* complex had one strain (0.5%), and *L. galeiforme* complex had one strain (0.5%). It is worth mentioning that all three species of the complex in this study were isolated from *Hylastes* sp. and its galleries, and only *L. wingfieldii* in the *L. clavigerum* complex was isolated from the *C. piceae*, Previous studies have shown that *Leptographium* spp. are highly represented among the fungi isolated from dying tree root tissues and bark beetles (Matusick et al., 2013), and that bark beetle can transmit spores of *Leptographium* spp. to injured or healthy root systems, causing root diseases and killing trees (Schweigkofler et al., 2005). The results of this study also demonstrate the associative relationship between *Hylastes* sp. and *Leptographium* and enriched the species of Leptographium-associated fungi.

We isolated *Ophiostoma ips* from the galleries of both species of bark beetles, of which 19 were related to *C*.*piceae* (9.2 %), and 9 were associated with *Hylastes* sp. (4.4 %). In China, *O. ips* is one of the most frequently isolated Ophiostomatales fungi, which has some specific symbiotic relationships with insects such as bark beetles, mites, and longicorn (Chang et al., 2017; Zhao et al., 2018), which was confirmed by our results. It is worth mentioning that whether the intersection of the two species of bark beetles in the ecological niche is a competitive or cooperative relationship needs to be explored by providing more data.

Meanwhile, this study’s fungi isolated from Sporothrix, Ceratocystiopsis, and Masuyamyces were all related to *Hylastes* sp. The association of the *Masuyamyces-Hylastes* relationship is the first time it has been found, indicating that many fungi related to *Hylastes* sp. have not been found.

An increasing number of studies have shown a symbiotic relationship between the Ophiostomatales fungi and the bark beetle. In this study, we identified the diversity of two Ophiostomatales fungi associated with two bark beetles in Liaoning, and determined the status of the new species in Ophiostomatales fungi. However, the current research on the associated fungi of the two species of bark beetles is mainly concentrated in Northeast China. We should continue to explore the diversity of Ophiostomatales fungi in different regions and further explore the relationship between fungi and bark beetles, which is of great significance for future research work.

## Acknowledgments

We thank all the forest staff who assisted us during the sampling process in Guangdong, Fujian, Guangxi, Guizhou, Shandong, Liaoning and Shannxi Provinces over the years, and we especially appreciate Professor J. Wang for his unwavering support and care for our laboratory-related work.

## Declarations

### Availability of data and material

The manuscript mentioned all the availability of data. Ex-type cultures of new species were deposited in the Culture Collection of South China Agricultural University (SCAU) and the China General Microbiological Culture Collection Center (CGMCC). The type herbariums were preserved in the Fungarium (HAMS), Institute of Microbiology, Chinese Academy of Sciences. DNA sequence data are available in Genebank (https://www.ncbi.nlm.nih.gov/nucleotide/), and taxonomic novelties are available in Mycobank (https://www.mycobank.org).

### Authors’ contributions

Conceptualization, Mingliang Yin; Data curation, Yutong Ran; Funding acquisition, Mingliang Yin; Formal analysis, Yutong Ran; Funding acquisition, Mingliang Yin; Investigation, Kun Liu, Minjie Chen, Yutong Ran and Congwang Liu; Methodology, Mingliang Yin; Project administration, Mingliang Yin; Resources, Yutong Ran, Kun Liu, Minjie Chen, and Congwang Liu; Software, Yutong Ran; Supervision, Mingliang Yin; Validation, Yutong Ran; Visualization, Yutong Ran; Writing – original draft, Yutong Ran; Writing – review & editing, Tong Lin and Mingliang Yin.

